# CEN-tools: An integrative platform to identify the ‘contexts’ of essential genes

**DOI:** 10.1101/2020.05.10.087668

**Authors:** Sumana Sharma, Cansu Dincer, Paula Weidemüller, Gavin J Wright, Evangelia Petsalaki

## Abstract

An emerging theme from large-scale genetic screens that identify genes essential for fitness of a cell, is that essentiality of a given gene is highly context-specific and depends on a number of genetic and environmental factors. Identification of such contexts could be the key to defining the function of the gene and also to develop novel therapeutic interventions. Here we present CEN-tools (<u>C</u>ontext-specific <u>E</u>ssentiality <u>N</u>etwork-tools), a website and an accompanying python package, in which users can interrogate the essentiality of a gene from large-scale genome-scale CRISPR screens in a number of biological contexts including tissue of origin, mutation profiles, expression levels, and drug response levels. We show that CEN-tools is suitable for both the systematic identification of genetic dependencies as well as for targeted queries into the dependencies of specific user-selected genes. The associations between genes and a given context within CEN-tools are represented as dependency networks (CENs) and we demonstrate the utility of these networks in elucidating novel gene functions. In addition, we integrate the dependency networks with existing protein-protein interaction networks to reveal context-dependent essential cellular pathways in cancer cells. Together, we demonstrate the applicability of CEN-tools in aiding the current efforts to define the human cellular dependency map.

## II. INTRODUCTION

A common approach to elucidate the function of a gene is to investigate the effect of its perturbation on a given biological process. Genetic perturbation experiments enable identification of genes that are essential for the survival and fitness of the cell. It is now widely accepted that the binary nature of gene essentiality as defined in classical genetics is too simplistic and gene essentiality is highly context-specific. There are genes, often termed as core fitness genes, which are required for core functioning and house-keeping of the cells and their knockout causes loss of fitness, in principle, in all cell types in all conditions. However, a large number of genes, termed context-specific essential genes, play important roles in providing fitness to a cell only in a particular genetic or an environmental context (Rancati *et al*, 2017). An important field in cancer research is the identification of genes whose loss is lethal to cells only in a specific context, as the genotype-specific vulnerabilities are excellent therapeutic targets for cancer cells carrying the specific genotype without affecting normal cells.

The recent advances in gene editing technology using the CRISPR/Cas9 knockout system, have enabled large-scale genome-wide screens to systematically perturb genes and rapidly identify those that are essential for proliferation and survival of cells (Tzelepis *et al*, 2016; Hart *et al*, 2015; Wang *et al*, 2015; Shalem *et al*, 2014). Initial studies on pooled essentiality CRISPR screens mainly focused on identification of therapeutically important context-specific vulnerabilities in different cancer types and mutational backgrounds (Steinhart *et al*, 2016; Tzelepis *et al*, 2016; Hart *et al*, 2015; Wang *et al*, 2017; Barbieri *et al*, 2017). In recent years, a number of studies have used the concept of co-essentiality-driven co-functionality—if essentiality profiles of two genes are correlated, the genes are likely to be involved in similar functions—to delineate novel gene functions using pooled CRISPR screens (Kim *et al*, 2019; Pan *et al*, 2018; Rauscher *et al*, 2018). Studies of this nature are possible because the number of cell lines that are being screened has increased to hundreds and thus it is viable to use the assumption that investigating the knockout fitness of genes across many different cell lines with different genetic backgrounds is comparable to studying different isogenic backgrounds of the same cell line. Together, the power of essentiality screens in systematic characterisation of cancer specific genetic-interaction maps of cellular processes and in the identification of therapeutically important genotype-specific vulnerabilities in cancer cells is now widely appreciated.

As the number of cell lines assayed for vulnerabilities has increased rapidly in the past few years, there has also been an increase in efforts to consolidate and standardise the essentiality screens performed in different laboratories. Recently, two major initiatives, DepMap (Meyers *et al*, 2017; Tsherniak *et al*, 2017) and Project Score (Behan *et al*, 2019), have performed essentiality screens in over 500 cell line models representing a wide range of tissue types using standard reagents and analysis pipelines. The PICKLES web server also provides a repository of all major essentiality screen initiatives, in which raw screening data are processed through a standard analysis pipeline (Lenoir *et al*, 2018). The availability of standardised essentiality screens combined with the efforts to characterise cell lines comprehensively in regards to the gene expression profiles, mutation profiles, drug responses, and copy number variations from initiatives such as Cell Model Passports (van der Meer *et al*, 2019) and Cancer Cell Line Encyclopedia (CCLE) (Ghandi *et al*, 2019; Meyers *et al*, 2017) now provide a premise for data integration for systematic studies that explores the genetic vulnerabilities in a wide range of contexts.

To aid the current efforts to democratise the large-scale screens and make them accessible to a broader scientific community, we here present CEN-tools. CEN-tools is an integrated database and set of computational tools to explore context-dependent gene essentiality from pooled CRISPR datasets, obtained from the two largest publicly available essentiality screening projects: the DepMap project and Project Score. CEN-tools offers an easily accessible web-interface with built-in statistical tools, to explore statistically significant associations between the essentiality of a given gene in a user-chosen set of cell lines and a predefined context (e.g. mutational background, expression levels, and tissues of origin). For advanced users, the python package implementation of CEN-tools also offers functions to interrogate bespoke contexts of choice to identify novel associations. We demonstrate that CEN-tools enables systematic studies to not only identify functional genetic interactions, but also to define the underlying context associated with the essentiality of a given gene. In addition, we showcase the applicability of CEN-tools as a new type of omics-database for integration with current -omics platforms to identify biologically relevant cancer-specific dependency networks.

## III. RESULTS

### CEN-tools identifies a robust set of core essential genes

Core-essential genes should, in principle, be essential in all cell types regardless of the mutational, tissue, or environmental background. To avoid setting arbitrary cut-offs as to the number of cell lines that define these genes, we developed a logistic-regression-based approach combined with clustering to categorise genes according to their essentiality probability profiles (Materials and Methods). The cluster of genes that exhibited high probability for being essential across all the cell lines was designated as the core-essential gene cluster. We separately analysed the Project Score (SANGER), and the DepMap project (BROAD) datasets and identified 650 genes from the SANGER dataset and 942 genes from the BROAD dataset, assigned into the core-essential gene cluster, with 519 overlapping genes between the two projects. Among these, 146 genes were previously annotated as core-essential genes by the ADaM (Adaptive Daisy Model) analysis tool, which is a semi-supervised algorithm recently used to identify novel core-fitness genes from essentiality screens in Project Score (Behan *et al*, 2019). We noticed that the core-analysis pipeline of CEN-tools was able to capture all but 20 genes previously identified by the ADaM pipeline suggesting that CEN-tools provides a robust platform to perform core-gene analysis (**Fig 1**). The 20 genes from the ADaM pipeline that were not captured among the high-confidence core-essential genes of CEN-tools were still identified in the SANGER dataset analysis of CEN-tools. Upon closer inspection, we observed a major discrepancy in the essentiality probabilities of these 20 genes between the two projects. Genes such as *LCE1E, MED31, PISD, UBB, ALG1L, HIST1H2BB* showed completely opposite essentiality profiles in the different projects (**Fig EV1**) suggesting differences in the gRNA efficiencies for those particular genes.

**Figure 1:**
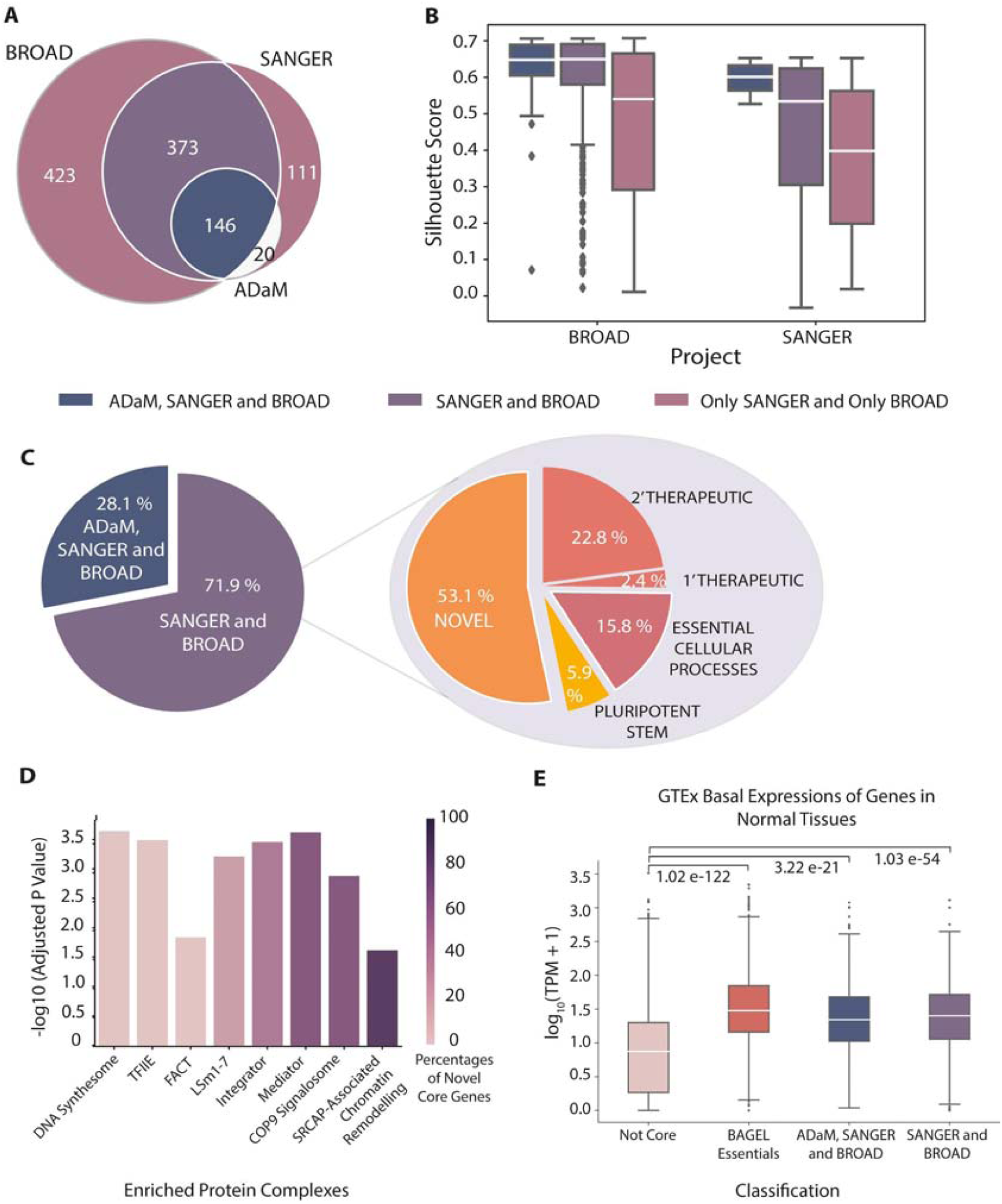
CEN-tools identifies core-essential genes involved in regular housekeeping functions of a cell. **(A)** Venn diagram for prediction of core essential genes by CEN-tools using both BROAD and SANGER projects, and ADaM core fitness genes (Behan *et al*, 2019). **(B)** Box-Plot for Silhouette Scores of core essential genes predicted by CEN-Tools and ADaM. **(C)** Pie chart for percentages of core essential genes predicted by both ADaM and CEN-Tools and only CEN-Tools by using both projects. The pie chart on the right panel represents the percentages of novel core genes from CEN-Tools, known core genes in pluripotent stem cells and core biological processes, and also therapeutically tractable genes annotated from the Project Score. “1’ and 2’ Therapeutic” refer to Project Score Class A and Class B drug targets respectively (Behan *et al*, 2019) **(D**) Bar Plot of protein-complex enrichment of core genes. Y-axis represents the significance of the enrichment; colours of the bars represent the percentages of the novel core genes in the complexes. **(E)** Box plot for the log value of the basal expression levels of core genes in BAGEL (Hart & Moffat, 2016), ADaM, CEN-Tools predictions, and non-essential BAGEL and non-core CEN-Tools genes.

To investigate how well the genes from each project were clustered, we next examined the Silhouette Score (Si-score) for each gene and observed that the genes that were designated into the core-essential cluster in both datasets and/or also by ADaM pipeline had higher Si-scores compared to the genes that were designated in only one of the two projects (**Fig 1B**). As a higher Si-score indicates that the object is well-matched to its own cluster, we defined the overlapping 519 genes from both projects to be the high-confidence set of core-essential genes from CEN-tools. Of the 373 newly identified core-essential genes, 59 could be assigned to known essential housekeeping complexes and processes, namely ribosomes, spliceosomes, proteasomes, DNA replication, and RNA polymerase, using the KEGG (Kanehisa *et al*, 2017) database. Another 22 genes were annotated to be essential for the fitness of human pluripotent stem cells (Ihry *et al*, 2019). To further filter genes that could be involved in fundamental cellular functions of a cancer but not a normal cell, we annotated 94 genes considered to be therapeutically tractable for multiple cancer types (Behan *et al*, 2019). This filtration step yielded a list of 198 ‘novel core’ genes presumably important for basic housekeeping of a cell (**Fig 1C**).

To identify the gene families enriched in the novel core gene lists of CEN-tools, we first explored the previously-annotated essential processes from the ADaM pipeline and were able to add new members to the pre-annotated enriched groups, such as additional members of the mediator complex *MED7, MED17* and *MED22* (**Table EV1**). We also independently performed protein-complex enrichment using the CORUM database (Giurgiu *et al*, 2019), which revealed enrichment in similar complexes relating to housekeeping functions of the cells such as the DNA synthesome complex, mediator complex, integrator complex, LSm1-7 complex, and the TFIIE complex. As a new core essential complex, we also identified enrichment in the COP9 signalosome complex (**Fig 1D**). As genes involved in core-functioning of cells are also expressed in cells of normal tissues, we next explored the expression of the newly annotated core-genes in normal human tissues from GTEX and observed that the basal expression of the newly annotated core genes was significantly higher than the genes that were not annotated as core-essential genes **(Fig 1E, Fig EV2)**. The complete set of high confidence core-essential gene lists is available to download from **Table EV2**. The “Essentiality profile” tab of the CEN-tools web-application provides essentiality profiles for individual genes for further browsing.

### CEN-tools enables rapid interrogation of contexts to identify gene-gene relationships

The ‘Context Analysis’ framework of CEN-tools calculates group-wise associations and correlations on a chosen set of cell lines to identify relationships between a given context and gene essentiality. To enable easy statistical comparisons, we preloaded a number of contexts on the CEN-tools website that are most likely to be used by the majority of researchers. These include tissue/cancer type-wide comparisons, essentiality correlations, correlation between essentiality and expression, essentiality driven by mutations in cancer driver genes, and correlation between drug responses and essentialities for predefined cell lines or user-chosen set of cell lines (see examples in **Fig EV3**).

Genes that are essential for a given tissue/cancer type were identified by testing if they had significantly higher essentiality compared to pancancer. Tissue specific dependencies, however, can be driven by a number of factors such as the underlying mutation that is enriched in the given tissue type or the level of expression of the gene in the particular tissue. Therefore, to get a better overview of different types of dependencies, we pre-calculated all possible associations for three main preloaded contexts (tissue/cancer, mutation, and expression), both in pancancer and within tissue/cancer type, and represent them in <u>C</u>ontext-specific <u>E</u>ssentiality <u>N</u>etworks or CENs. Each edge in this network represents the type of association, using which we annotated the underlying contexts. To demonstrate the value of CENs, we collected the co-essentiality networks from the PICKLES database (Lenoir *et al*, 2018) and extracted the corresponding genes from our CENs for direct comparison. On the *BRAF* CEN, for example, we could identify components of the co-essentiality networks with the mutational links of *BRAF* to *MAPK1, MAP2K1, PEA15, NFATC2*, and *DUSP4* **(Fig 2A, Fig EV4)**. However, *SOX10, MITF*, and *ZEB2* were not linked to BRAF itself but were instead associated to *BRAF* via their expression in the skin. *SOX10* and *MITF* additionally contained high-confidence self-loop edges of expression-essentiality correlation suggesting that those dependencies are not directly related to *BRAF* mutational status but rather through their expression status in skin, consistent with the lineage specification roles these genes play in skin tissue regardless of the mutational background (Harris *et al*, 2013).

**Figure 2:**
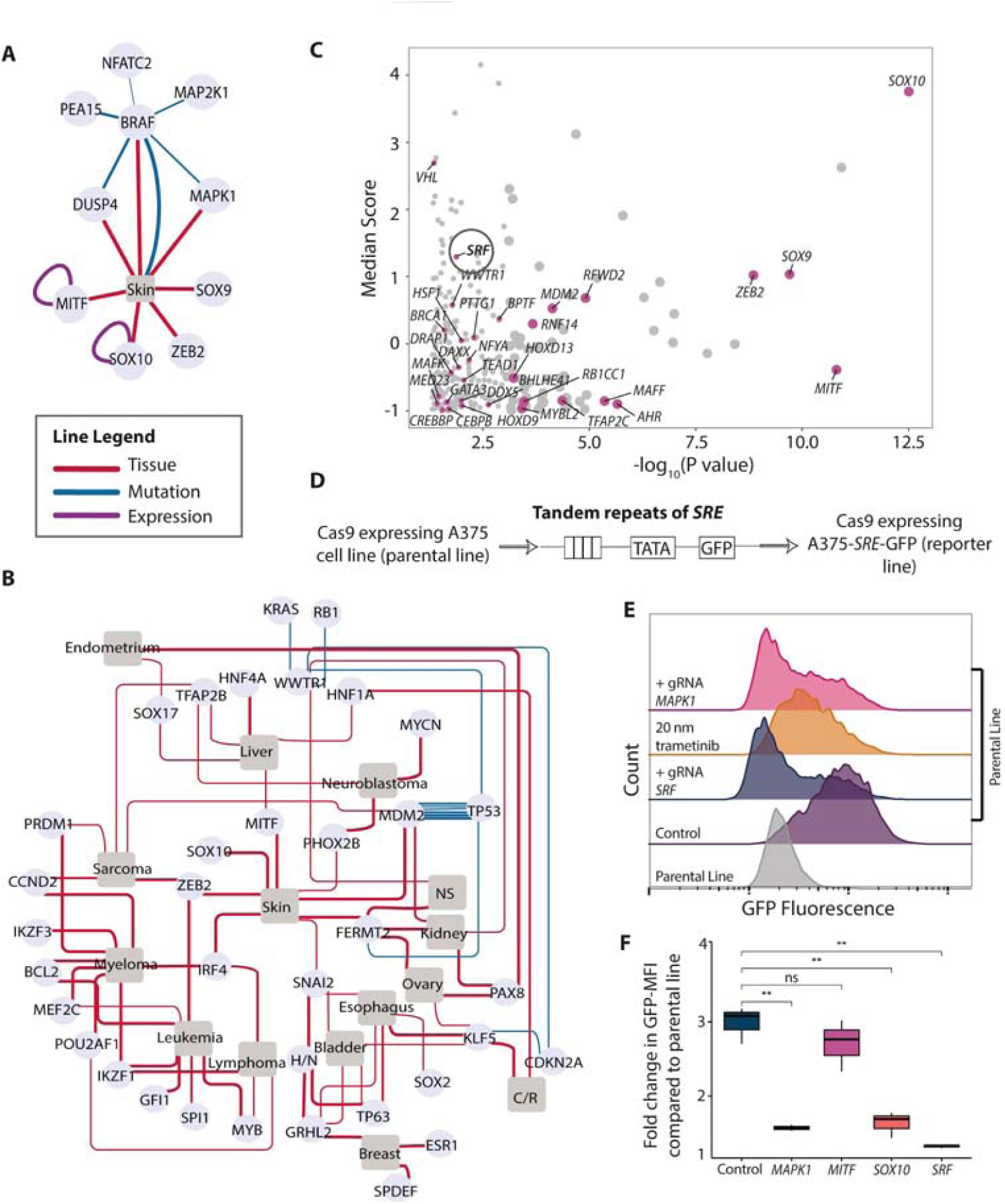
Interrogation of contexts from CEN-tools identify novel gene-gene relationships. **(A)** An example of CEN from CEN-tools-in this example, components of the *BRAF* co-essential genes from PICKLES were extracted from CEN-tools. **(B)** CENs for genes that have restricted expression and essentiality in different tissue types. Majority of the genes in this network are TFs important for lineage specification of a cell line to a particular tissue type. The width of the line in (A) and (B) denotes the confidence of association. **(C)** Genes with high essentiality in skin cell type compared to pancancer. Only the genes encoding for the transcription factors are labelled. The median score refers to the median of the scaled-Bayes essentiality score for the 32 skin cell lines from the DepMap project. **(D)** Schematic of the Cignal^®^ lentiviral reporter construct for assessing the activity of the serum response factor (SRF) transcription factor. The construct expresses GFP under a control of a basal promoter element (TATA box) together with a multiple tandem repeats of serum response element (SRE). This construct was used to generate a reporter Cas9 expressing A375 cell line for SRF activity **(E)** Representative histograms depicting the GFP expression from the parental and the reporter line under different perturbations. For gRNA transductions, polyclonal lines were used to assess the GFP expression 6 days post-transduction. GFP expressions of trametinib treated cells were measured 2 days post treatment. **(F)** Boxplot depicting the GFP fold change distribution compared to from parental line GFP distribution from three independent gRNA transductions targeting the indicated genes. Polyclonal knockout lines were used for quantification. Representative raw-FACS plots from one of the three replicates are also shown in **Fig EV5C.**

In addition, the CENs were particularly useful in navigating through specific types of dependencies. For example, genes required for tissue differentiation into a particular lineage often have restricted expression, and given their cellular function, they should have high essentiality in the given tissue. Using CEN-tools, we isolated such genes (**Fig 2B**) and revealed a sub-network consisting of a number of transcription factors that are known to control tissue differentiation into a specific lineage such as *SOX10* and *MITF* in skin (Harris *et al*, 2013), *PAX8* in ovary, kidney, and endometrium ((Cheung *et al*, 2011; Tong *et al*, 2011; Grote *et al*, 2006), *MYCN* in neuroblastoma (Huang & Weiss, 2013), and *HNF4A* in liver (DeLaForest *et al*, 2011). The transcription factor *TP63* was highly expressed and essential in cell lines derived from head and neck, oesophagus, and bladder cancers, consistent with it being a known regulator of squamous epithelium lineage (Network & The Cancer Genome Atlas Research Network, 2012). Cell lines derived from cancers of blood cells are known to have distinct lineage specification genes and we also observed multiple specific lineage markers such as *MEF2C, IKZF3, IRF4, SPI1, POU2AF1, BCL2* (Behan *et al*, 2019; **Fig 2B**). *ZEB2* was associated with skin, haematopoietic and lymphoid, and soft tissue with a high statistical confidence, which is consistent with the mesenchymal origin of the cell lines from these tissue origins (De Craene & Berx, 2013). This subnetwork also revealed genes that are not necessarily lineage restricted but have an expression to essentiality relationship because of an underlying enriched mutational background. For example, the essentiality of *MDM2* in multiple tissue types was higher in cells with wild type (WT) *TP53.* This is an observation often made in pooled CRISPR screens, in which deletion of a repressor of a tumour suppressor gene in a cell line harbouring the WT tumour suppressor gene leads to very high proliferation causing identification of the repressor as an essential gene (Hart *et al*, 2015). Together, these examples illustrate how networks from CEN-tools can be utilised to systematically characterise specific types of expression-related dependencies from large-scale CRISPR screens.

### CEN-tools reveals a skin-specific link between the SOX10 transcription factor and SRF activity

We next examined if tissue/cancer specific networks could be explored in a similar manner to identify context-specific gene function. As a case study, we selected the skin tissue and extracted all transcription factors (TFs) that were directly linked to the skin tissue as TFs are most likely to play a central role in controlling tissue specific gene expression. The skin TF CEN revealed a number of lineage-specific markers such as *SOX10, MITF*, and *ZEB2* but also a number of other TFs whose expression is not restricted to the skin cell type (**Fig 2C, Fig EV5A**). The essentiality of a number of these TFs were associated with the mutational background of skin such as *TFAP2C, BPTF*, and *AHR* to *BRAF* and *TEAD1* to *NRAS*. Of the TFs whose essentiality was not associated with enriched mutations of skin, *SRF* was of particular interest to us as it is known to be activated through the MAPK pathway, which in turn is known to be dysregulated in melanoma cell lines harbouring the *BRAFV600E* activating mutation.

To further investigate the role of *SRF* in the context of melanoma, we opted to utilise the A375 melanoma cell line harbouring the BRAF activating mutation. To this end, we generated a clonal Cas9 expressing reporter version of the A375 cell line that contained an expression cassette for GFP driven by a serum response element (SRE) promoter containing eight binding sites for SRF (**Fig 2D**). We noticed that the reporter cell line constitutively expressed GFP when grown in media containing serum, which suggested that SRF was constitutively active in these cell lines. To ensure that the expression of GFP was as specific to the activity of *SRF*, we transduced the reporter cells with a single gRNA targeting *SRF*, which abolished the expression of GFP (**Fig 2E)**. To test if the activating mutation in *BRAF* and the consequent upstream hyperactive MAPK pathway acted on downstream *SRF* on these cell lines, we targeted components upstream of *SRF* with trametinib, which is an inhibitor of MAP2K1/2 kinase and also transduced cells with single gRNA targeting *MAPK1.* Both of these treatments led to a decrease in the GFP signal indicating that the activity of *SRF* in these cell lines was specific to the MAPK pathway (**Fig 2E**). While the dysregulated MAPK appeared to act directly on the activity of *SRF*, the essentiality of *SRF* in skin tissue was not related to the *BRAF* mutational status of the cells (**Fig EV5B**). We hence tested the effect of perturbing TFs with skin restricted expression on the activity of *SRF*. While targeting *MITF* with a single gRNA did not have an effect on *SRF* activity, we noticed a significant decrease in GFP expression when *SOX10* was targeted, indicating that the activity of *SRF* was related to the expression of *SOX10* (**Fig 2F, Fig EV5C**).

### CEN-tools uncovers essential cellular processes in cancer

Identification of mutation-dependent vulnerabilities are crucial for designing drugs that target cancer cells bearing such vulnerabilities without affecting the normal cells. To explore these vulnerabilities, we focused on the mutational associations identified in our CENs. As gain-in-function mutations in oncogenes are often associated with an increase in dependence of the cell lines harbouring the mutation, we first extracted self-loop edges connected by mutation driven essentialities in a pancancer comparison. The occurrences of oncogenic mutations, however, can be tissue-specific, so we additionally extracted edges corresponding to mutational association between oncogenes and tissue/cancer types, which was generated by comparing the essentiality of the given cancer driver in the context of its mutation within a given tissue/cancer type. Among the most significant associations were the pancancer mutational association in genes such as *BRAF, KRAS, NRAS, HRAS, CTNNB1, PIK3CA, EZH2* in at least one of the two projects (**Fig EV6**). Within group A associations (statistical tests with six or more cell lines/group), a number of these genes were associated with tissues such as *BRAF* with Skin, *PIK3CA*-Breast and ovary, *KRAS* with pancreas, oesophagus, colon/rectum, and lung and *NRAS* with skin and hematopoietic and lymphoid, indicating the tissues in which these mutations are most relevant in.

CRISPR-screens performed on cancer cells with a particular genetic background are also perfectly suited to identify synthetic lethal interactions. We examined if mutational networks of CEN-tools could be used as a guide to identify synthetic lethal interactions in tissues with a specific mutational background. As a case study, we investigated the edges corresponding to increased essentialities in NRAS mutational skin tissue background. NRAS mutations constitute 15-20% of all melanomas and are the most important sub-group of BRAF wild-type melanomas, yet therapeutic options for NRAS mutant melanoma are still limited (Muñoz-Couselo *et al*, 2017). It is known that NRAS mutant cancer cell lines rely on signalling through *CRAF* (*RAF1*) and *SHOC2* (Dumaz *et al*, 2006; Jones *et al*, 2019; Kaplan *et al*, 2012) and we could identify this association as a highly significant association in CEN-tools (**Fig 3A**). To identify the cellular context of these dependencies, we opted to integrate the dependencies from the relevant CEN with a protein-protein interaction network to identify affected cellular mechanisms. The integrated dependency-interaction network (**Fig 3B**) revealed a cluster of highly connected components with significant enrichment in a number of pathways including the Ras-signalling pathway, the focal adhesion pathway and the m-TOR pathway (**Fig 3C**). An important protein of the Ras-signalling pathway identified within these clusters was the receptor tyrosine kinase IGF1R. A closer look revealed that the essentiality of *IGF1R* was significantly higher in *NRAS* mutant cell lines compared to *NRAS* WT background and also generally in *BRAF* WT melanoma cell lines, which is mainly composed of *NRAS* and *HRAS* mutants, compared to the BRAF V600E mutants (**Fig 3D**). In addition to *IGF1R* itself, genes encoding the FURIN protease, which is required for surface presentation of *IGF1R* (Kavran *et al*, 2014) also had a higher essentiality in *NRAS* mutant melanoma cell lines compared cells with WT *NRAS* (**Fig EV7**). The role of *IGF1R* in mediating acquired resistance of *BRAF* mutant melanoma cells to *BRAF* inhibitors is well-studied (Corcoran *et al*, 2011; Villanueva *et al*, 2010); however, the role of *IGF1R* in *NRAS* mutant melanoma is not completely understood and further work will be required to establish the precise manner in which *NRAS* mutant melanoma cell lines are dependent on *IGF1R*. Additionally, the identification of genes encoding proteins such as *ITGAV, RHOA*, and *RAC1* suggest a potential role of cellular components controlling cell adhesion and cytoskeletal organization in NRAS mutant melanoma. Together, this case study suggested that integrating the protein-protein networks with CENs provides a powerful means to identify potential synthetic lethal interactions and at the same time represent the genes from dependency networks into interaction modules with enriched cellular functions for better elucidation of cell essential biological processes.

**Figure 3:**
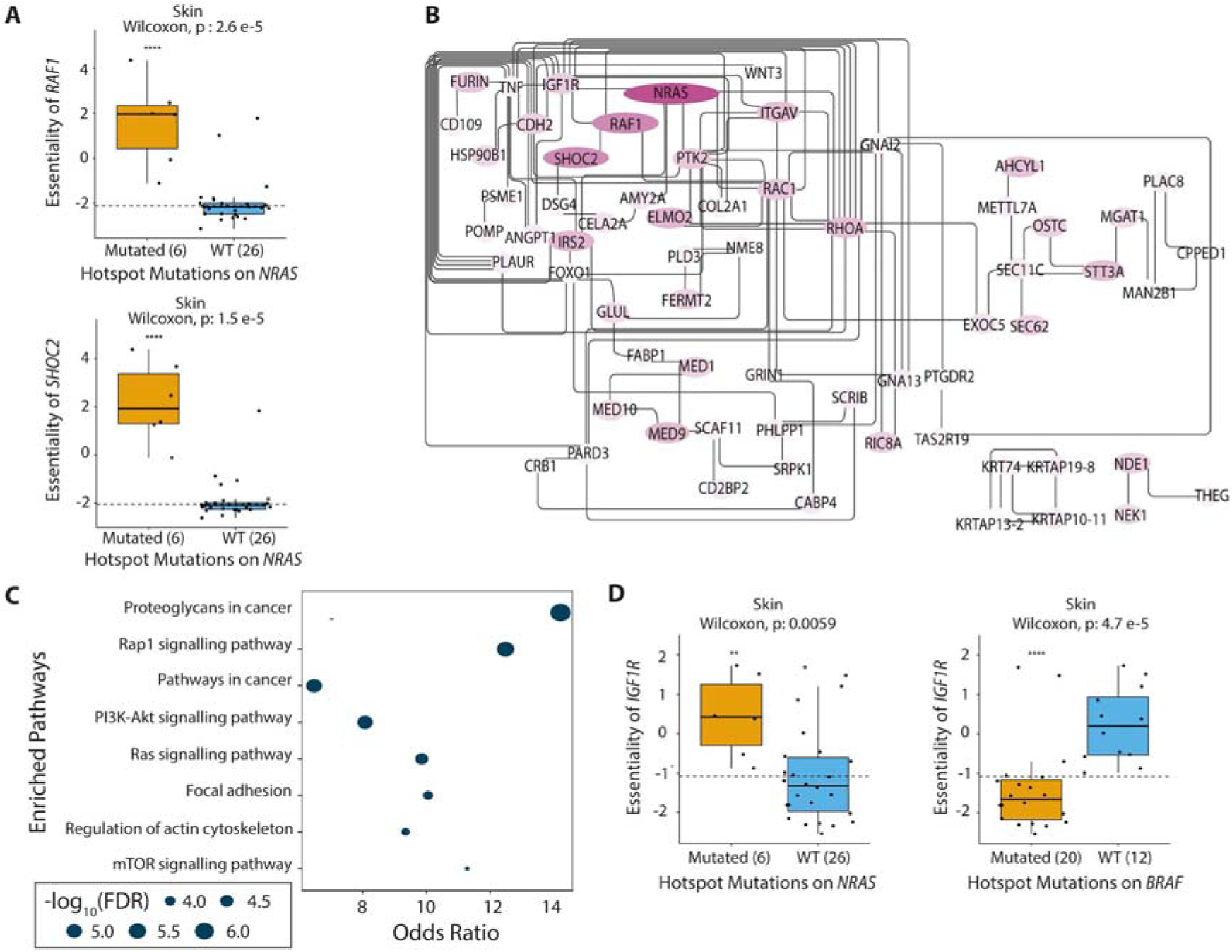
CENs can identify cell essential processes for NRAS mutant melanoma cell lines. **(A)** The essentialities of *RAF1* and *SHOC2* are higher in NRAS mutant melanoma cell lines compared to NRAS WT melanoma cell lines. **(B)** Protein-protein interaction network of essential components of NRAS mutant melanoma cell lines. All nodes in the network are genes whose essentiality is significantly higher in NRAS mutant melanoma compared to NRAS WT melanoma cell lines. The colour intensity represents the median-change in essential scores of NRAS melanoma cell lines compared to that of NRAS-WT cell lines. **(C)** Enriched pathways in the network in **(B). (D)** Boxplots depicting the higher essentiality of *IGF1R* in NRAS mutant melanoma cell lines compared to non-mutant melanoma (left panel) and lower essentiality of *IGF1R* in *BRAF* mutant melanoma cell lines compared to *BRAF* WT melanoma lines.

## IV. DISCUSSION

Loss-of-function genetic screens are a powerful means to identify cellular vulnerabilities and such screens are now available for a large number of genomically and transcriptomically characterised cancer cell lines. Despite their obvious value to understanding gene functions these screens are under-represented in omics-integrative systems biology approaches. This is largely due to the fact that the ‘essentiality’ of a given gene is highly context-dependent, making the interpretation of essentiality from a genetic screen non-trivial and largely inaccessible to non-experts.

To address this, here we present CEN-tools, a website and accompanying python package that integrates these data. CEN-tools allows the easy navigation of the major publicly available large-scale genetic screens to identify associations between a given context and the essentiality of a gene. For this purpose, it provides a suite of statistical tools as well as integration with a number of contexts from publicly available datasets, such as gene expression, mutation background, and drug sensitivity profiles of cell lines. Additionally, the ‘Cell Line Selection’ tool within CEN-tools further enables users to restrict their analysis to their context of choice, regardless of the pre-loaded contexts, and make comparisons for interesting contexts such as paralog dependencies within a cell, essentialities driven by gene amplifications, and essentialities associated with defects in DNA-repair mechanisms (examples in **Fig EV8**). Further flexibility to investigate contexts of choice is also available through allowing the upload of a custom list of interesting cell line IDs. The python pipeline of CEN-tools also enables systematic studies using multiple genes as queries, a feature we plan to include in the future version of the website. An application for this, for example, could be to query dependencies in the context of a specific cancer subtype signature.

As a basis for CEN-tools, we identified high confidence core-essential genes, through an analysis that combined the essentiality profiles of genes from the two projects. Our core-gene analysis captured almost all genes identified by ADaM (Behan *et al*, 2019), and identified 198 new core essential genes involved mainly in previously known-to-be-essential cellular processes, but also added an additional process of COP9 signalosome complex to these. The core-analysis from CEN-tools could also capture all but 11 genes from the very recent core-gene analysis from Dede *et al. (Dede et al, 2020)*, which used the same datasets as in this study (**Fig EV9**). As the essentiality measured from genetic screens not only captures genes whose loss causes cell death, but also genes whose loss results in slower proliferation of the cells, the day the experiments were performed will affect the identification of the genes essential for proliferation. This perhaps explains the higher number of core-genes identified in the BROAD datasets compared to the SANGER study (21 days instead of 14 days). By directly comparing the two studies we also identified cases in which the gRNAs’ efficacy in the two projects varied considerably. Genes like *UBB, HIST1H2BB*, which encode for Ubiquitin B and a histone protein are very likely to be core-essential but were not identified as such in the BROAD study which suggests that the gRNAs used for these particular genes in the BROAD study were of lower efficacy. Our high confidence core-genes were defined as those that overlapped in both projects and hence this core-essential gene list from CEN-tools is representative of genes in which gRNA efficacy for a given gene in both studies were comparable. However, users of CEN-tools can easily navigate the essentiality distributions for both projects to identify both the best timeframe and library for their specific genes of interest, when designing knockout experiments.

Using the CEN-tools for groupwise testing for significant association, we identified many previously described molecular markers that are associated with essentiality, including, but not limited to mutational dependence on major cancer drivers and tissue specific lineage dependency markers (Behan *et al*, 2019; Tsherniak *et al*, 2017). We further used CEN-tools to discover and experimentally validate an association between the skin-specific gene *SOX10* and the *SRF* transcription factor downstream of MAPK signalling in malignant melanoma. Additionally, we showed that integration of CENs with existing protein-protein interaction networks provides a powerful way to map dependencies in the context of cellular function. In the example of NRAS melanoma, in addition to identifying the expected Ras-signalling pathway, we also identified a potential role of cellular cytoskeletal processes, a pathway that is not entirely evident by only exploring individual dependencies. While these results require validations through further experiments, the results here suggest that our approach of CEN-PPI integration is both viable and novel, not yet commonly applied in the field of systems biology, to identify cellular pathway dependencies. Approaches of this nature could be refined in the future as more context-specific PPIs (e.g patient-specific PPI interactions) become available to ultimately aid designing better therapeutics.

It should be noted that as the current version of CEN-tools tests group-wise associations for only the given context without taking into consideration other co-occurring contexts, there is a risk of observing confounding results if one were to only perform a single test in isolation. We thus recommend considering associations in the context of their CENs, through the network view of CEN-tools, as these will be able to point to some of the confounders, at least for the contexts whose associations have been pre-calculated. All systematic analysis within CEN-tools have been performed separately for the two projects to avoid variations due to institute origins of the cell lines. The cell line selector tool could be used for further exploration of possible confounders when interpreting associations, as it is equipped with additional information about the genetic and transcriptomic makeup of each cell line. Future versions of CEN-tools, and as more data becomes available, will integrate an analysis for confounding factors more directly. An option for this could be identifying associations between gene essentialities and a given context using a mixed effect linear model while considering defined set of contexts as covariates, an approach that has been used very recently for the identification of drug-gene associations from essentiality screens (Gonçalves *et al*, 2020). Currently, to aid the users in interpreting the statistical associations we have included a number of confidence annotations. For example, all analyses are provided at two levels of confidence: Group A analyses include a sufficient number of cell lines for confident statistical associations, and Group B should be used with caution as it is restricted by a low number of available cell lines in the datasets. All statistical associations of CENs are also annotated with low to high confidence levels depending on the p-value or correlation scores, enabling the users themselves to filter the associations depending on their requirements.

In summary we have developed a platform that can be used to explore the dependency of a given gene in a given context, which is key for elucidating the molecular function of a gene. The flexible and modular nature of CEN-tools allows for future integration with several other interesting data types, as they become available in sufficient numbers, such as drug synergy data and other context specific information, such as protein expression, post translational modifications and signalling signatures in specific contexts. Thus, we expect that CEN-tools will make essentiality screening data widely accessible to researchers, and will facilitate its integration with orthogonal datasets, to perform systematic studies to identify cancer dependencies.

## Author contributions

SS and EP conceived the study. SS, CD led and designed the study and performed all analyses for CEN-tools. SS and PW designed and implemented the web server. CD created the python-based advanced CEN-tools suite with input from SS. SS and CD wrote the manuscript with input from EP and PW. SS and EP co-supervised CD and PW. EP supervised SS and provided input and feedback on all aspects of the study. GJW and EP supervised SS on the experimental part of the study.

## Acknowledgements

SS, CD, PW and EP and the manuscript publication costs are funded by EMBL-EBI core funding. GJW and experiments were funded by Wellcome Trust (grant 206194). We would like to thank the Wellcome Sanger Institute flow cytometry Core facility for help with flow cytometry experiments; James Klatzow for help with core-gene analysis; Ijaz Ahmad for help with setting up the web server and the members of the Petsalaki group, Toby Gurran and Mathew Garnett for critical reading of the manuscript and testing of the CEN-tools web server.

## V. MATERIALS AND METHODS

### Data collection and curation

Essentiality screens for cell lines and the associated information on expression, CNV, drug response, and mutation were obtained from publicly available databases **(Table 1)**. In addition, a number of other datasets were used in annotation of genes and their functions (**Table 1, Table EV3**). Essentiality scores from Project score and DepMap projects were retrieved from Project Score, in which the same pre-processing pipeline was applied. To make comparison between the two projects feasible, all systematic analysis was restricted to the genes targeted in both projects (16819).

**Table 1.**
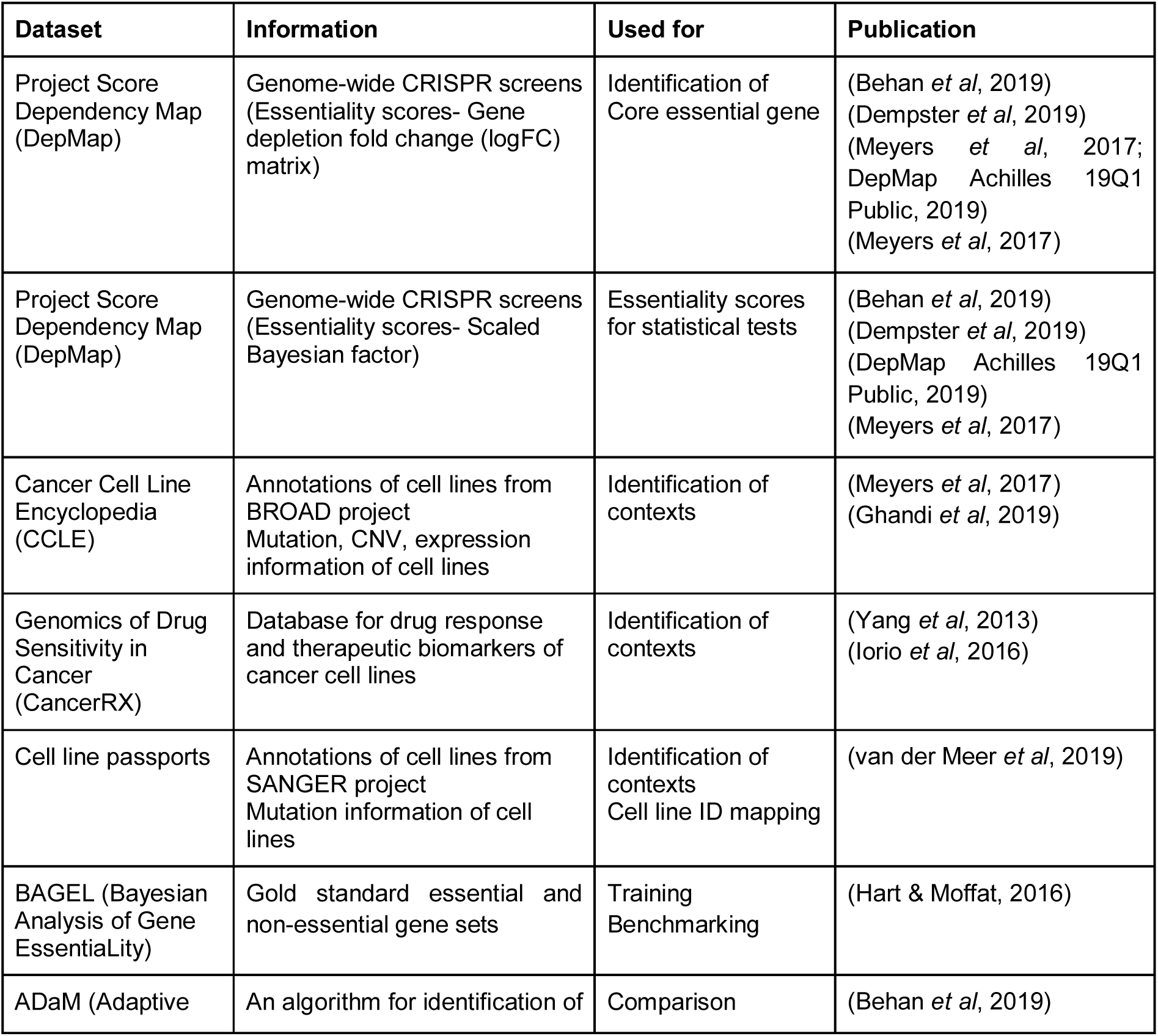
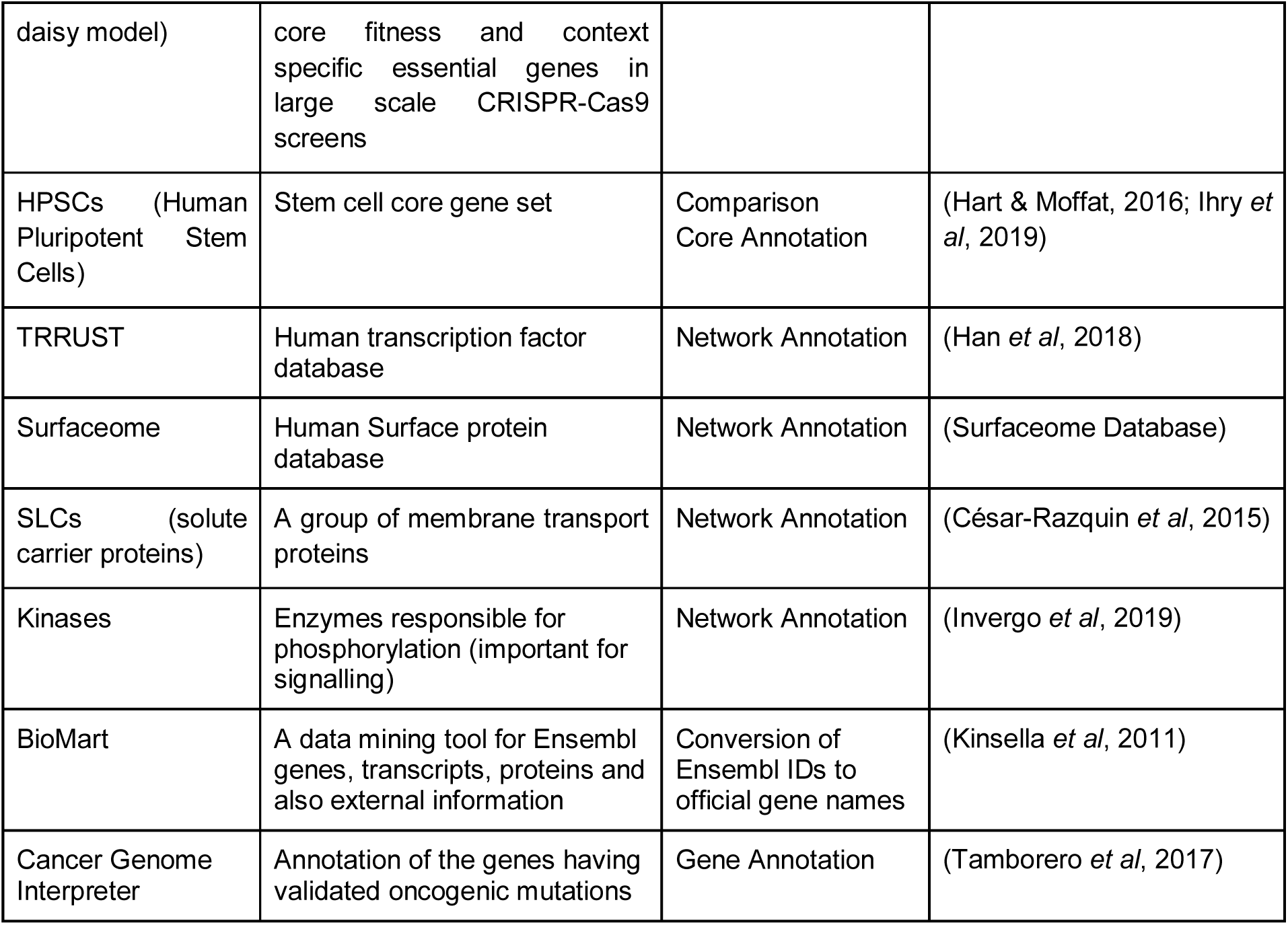
Datasets used in this study.

### Identification of core essential genes

To identify core genes that are essential for all cell types, we used the normalised log-fold-change (logFC) matrix calculated using the MAGeCK (Li *et al*, 2014) pipeline post CRISPcleanR (Iorio *et al*, 2018) correction from the project score analysis pipeline. Rather than using strict cut-offs to identify essential genes from each dataset, we opted to convert the logFC values into essentiality probabilities and compare the probability of each gene being essential from the two independent datasets to identify core essential genes. To this end, we applied the following three steps: (i) Training a logistic regression function that can separate genes as essential and non-essential inside each cell line, (ii) Extracting probability distributions of the predictions for being essential of each gene across cell lines and finding the patterns of the distributions, and (iii) Clustering the essentiality patterns using KMeans and defining core essential and non-essential gene clusters as shown in **Fig EV10A**.

First, we separately trained logistic regression models with gold-standard reference BAGEL essential and non-essential gene lists from R implementation of BAGEL (Hart & Moffat, 2016). For each project, we separately applied these models on the remaining genes and assigned probabilities of each gene being essential in each cell line (ROC and PR Curves in **Fig EV10B**). Then, to calculate the patterns of probabilities for a gene being essential across cell lines, we first sorted the essentiality probabilities in an increasing order and converted the continuous probability distribution into 20 bins of discrete frequency of probability distributions. This generated a numeric matrix of probability frequencies for each gene being essential. Finally, we used these matrices as input for unsupervised k-means clustering. Using the silhouette method, we identified the optimum number of clusters to be 4. The four major clusters of probabilities represented the different nature of gene essentialities as shown in **Fig EV10C**. We identified Cluster 1 as the cluster that best represented core-essential genes as the genes in this cluster had probability distribution skewed to 1 as a peak. Cluster 4 genes were the opposite of Cluster 1. The remaining two clusters indicated context specific essential genes as their probability distribution revealed essentiality fluctuations across the cell types.

### Contexts interrogated in CEN-tools and representation of the dependency networks (CENs)

The pre-defined contexts in CEN-tools were split into either “discrete” or “continuous context” and were broadly classified into three major groups:

1. Tissue/cancer (discrete context): Tissue refers to the tissue of origin of the cell lines. Cancer refers to the further classification of the cell lines into the cancer type within each tissue. Supplementary table 2 shows the different tissues and cancer types available in the two different databases and the relationship between the cancer types with the tissue of origin. For statistical testing, the chosen tissue was used as the “test” group.
2. Mutation (discrete context): For mutation analysis, two different mutational annotations obtained from CCLE database were used. Both annotations were obtained from CCLE website (Meyers *et al*, 2017; Ghandi *et al*, 2019). In all two group statistical testing, for a given gene, the cell lines with the mutated gene were used as the “test” group and those with non-mutated gene were used as the “control” group.
  a. CCLE:Hotspot mutation: Pre-annotated commonly occurring hotspot mutations in 75 genes were used in the analysis. Specific protein changes for the hotspot mutations used in this study are detailed in **Table EV4A**.
  b. CCLE:Oncogenic mutation: A less stringent mutational analysis was performed by using *any* mutation in an oncogene. A total of 76 oncogenes were annotated (**Table EV4B**).
3. Expression (continuous context): For expression analysis, cell line expression values were obtained from the CCLE database as normalised FPKM values (Ghandi *et al*, 2019; Meyers *et al*, 2017).
4. Drug response: Drug response for the cell lines were obtained as normalised Z-scores from the CancerRxGene (Yang *et al*, 2013; Iorio *et al*, 2016).

To identify the contexts that were associated with essentiality, we used the scaled Bayes-factor (BF) essentiality matrix from Project Score (Behan *et al*, 2019). In this matrix, a score < 0 indicates a statistically significant effect on cell fitness. Contexts were first separated into continuous or discrete type contexts.

For discrete contexts, cell lines were separated into test and control groups depending on whether they fulfilled the criteria of the context or not. The groups were separated either from pancancer or within a specific tissue/cancer type. We opted for non-parametric tests for group-wise testing mainly because the scale BF essentiality scores were not normally distributed and also in some cases the number of samples in each group were fewer than 20. For the statistical tests, Kruskal Wallis and Mann Whitney U (two-samples Wilcoxon) tests were used with default parameters (*SciPy* v.1.4.1 package (Virtanen *et al*, 2020) in Python v. 3.7.4 and *ggpubr* package (Kassambara) in R). For all statistical tests, the number of samples in each group was at least 3. For further confidence annotation, associations were divided into group A type association, in which there were more than five samples per group, and group B associations, in which there were only 3-5 samples per comparison group.

For continuous contexts, pearson correlation was used (*SciPy* v.1.4.1 package (Virtanen *et al*, 2020) in Python v. 3.7.4 and *ggpubr* package (Kassambara) in R). Correlation calculations are provided between the selected group and pancancer as well as within the relevant tissue/cancer type. Correlation tests required at least 5 samples. In the website version of CEN-tools, correlations were performed using *ggpubr* package (Kassambara) in R.

The statistical tests yielded a large number of associations between genes with the pre-defined contexts. To extract meaningful relationships, we iteratively searched pre-defined contexts in all genes in both projects and used a network visualisation approach to represent all statistical relationships.

### Integration of CEN with human interactome

To elucidate the cellular essential processes of the *NRAS* mutant skin cancer cells, we integrated the *NRAS* mutant skin CEN, comprising the statistical associations between the gene dependencies, with the STRING protein interaction network, comprising biological associations in the human cells. To do this, first we extracted the *NRAS* mutant skin CEN from whole dependency network, filtered by: project=BROAD, tissue=NRAS mutated skin tissue cell lines, effectors=mutation and co-mutation, and direction=enhancing the essentialities of the affected genes in the presence of *NRAS* mutation. Second, we obtained the human interactome from the STRING database (Szklarczyk *et al*, 2019) and constructed a network object (*networkx* package (Proceedings of the Python in Science Conference (SciPy): Exploring Network Structure, Dynamics, and Function using NetworkX) in Python v. 3.7.4). Then, to obtain the biological associations between the dependent genes, we calculated the shortest path between genes connected by each edge in CEN, iteratively. The genes in all the shortest paths between CEN edges in PPI were used to extract the subgraph of the interactome as an integrated network (**Table EV5A, Table EV5B** is the network file and node attribute file, respectively).

Since we only focused on the dependent genes and their signalling routes, we removed all the nodes that were absent from the CEN and all the edges absent from the interactome. In order to make the network stringent, we removed dependent genes having median essentiality below than or equal to −1.5 in *NRAS* mutant skin cell lines, edges having STRING interaction score below 400, singleton nodes, and nodes having only one neighbour. Pathway enrichment on the finalised nodes was performed using Enrichr package (Kuleshov *et al*, 2016) in R.

### Cell line selection tool

To allow users to choose specific cell lines to explore using CEN-tools, the Shiny app implementation offers the upload of a list of cell lines. For a more explorative and interactive approach, the user has also the option to custom select cell lines using our “Interactive Cell Line Selector”. Using simple drop-down menus, the user can restrict the choice of cell lines based on mutations, copy number variations (CNV), growth property, microsatellite stability, tissue and cancer subtype. Only cell lines for which we had essentiality data, mutation information and expression data are available for selection.

Cell lines adhering to the made selection will be presented in a table and highlighted in a T-distributed Stochastic Neighbour Embedding (t-SNE) plot for further interactive selection. In the t-SNE plot, each cell line is represented by a dot. Cell lines with similar gene expression profiles will be closer together. To generate the t-SNE plot, the gene expression matrix of each project (BROAD and SANGER) containing FPKM values with genes as columns and cell lines as rows was reduced to two dimensions using the *Rtsne* package (van der Maaten, 2014; Maaten & Hinton, 2008) in R (version: 0.15). Default parameters were used except perplexity was set to 50 and the number of maximal iterations was changed to 500. The t-SNE plot was generated with the R *ggplot2* package using the scatterplot function *geom_point* (Wickham, 2009) (version: 3.2.1) with the two t-SNE dimensions as x and y axis, respectively. Cell lines with similar expression profiles are close together in the two t-SNE dimensions. The user can select cell lines interactively by either clicking on the dots of interest or by drawing a window over a number of cell lines and hitting the “Select points” button. To support the decision, the dots can be highlighted by tissue of origin or subtype. The choice of selectable dots can be restricted by using the above-described drop-down menus. As soon as the user is satisfied, the selected cell lines can be submitted for further analysis to CEN-tools.

Advanced python tool users of CEN-tools are also able to restrict the analysis to their choice of cell lines. For detailed guidance on the python tool-refer to the CEN-tool python documentation.

### Experimental methods

#### A375 cell culture

Cas9-expressing A375 cell line was obtained from the Sanger Institute Cancer Cell Line Panel (https://cancer.sanger.ac.uk/cell_lines). The cell line was cultured in DMEM/F12 media (Life Technologies) supplemented with 10% heat-inactivated (50°C for 20 min) FBS, 20 µg/mL blasticidin, and penicillin–streptomycin at 37°C with 5% CO_2_. Logarithmic growth phase of the cells was maintained by passaging the cells every 2-3 days. Cells were tested and found to be mycoplasma free.

### Lentiviral production and transductions

All lentiviruses were produced and all transductions were performed as described before (Sharma & Wright). All gene-specific gRNAs were obtained from **Sanger Whole Genome CRISPR Arrayed Libraries** (Metzakopian *et al*, 2017) and lentiviruses were produced in HEK293-FT cells using the Addgene lentiviral packaging mix. Polybrene (8 µg/mL) was added to the cells prior to the transduction of A375 cells using the ‘spinoculation’ procedure.

### Generation of the reporter A375 cell line

The reporter line of A375 was generated by transducing Cas9 expressing A375 cell line with a commercial reporter construct for SRE activity (Cignal Lenti SRE Reporter (GFP) Kit: CLS-010G). To establish a clonal reporter line, eight days post transduction, cells were individually sorted into 96-well plates using FACS (MoFlo XDP). The clonal cells were further cultured for 3 weeks and 48 individual clones were tested for their expression of GFP. Five clones exhibiting different levels of GFP were selected for expansion. The selected clone exhibited the highest constitutive expression of GFP when grown in culture supplemented with 10% serum and the highest loss of GFP signal when transduced with a single gRNA targeting *SRF*.

### Flow cytometry

Drug treatments and gRNA transductions were performed on six-well culture plates with 1×10^6^ cells/well. Treated cells were detached from culture plates using EDTA, washed twice with PBS supplemented with divalent ions and analysed using flow cytometry. All flow cytometry was performed on a Cytoflex flow cytometer. 10,000 events per sample was acquired for each sample. Live cells were gated using forward and side scatter. GFP was excited at a wavelength of 488 nm and emission detected using a 530/30 band pass filter; BFP was excited at 405 nm and the emission detected using a 450/50 band pass filter. Analysis was performed using FlowJo software (Treestar).

## VI. DATA AND SOFTWARE AVAILABILITY

All code used for the core essentiality analysis and the CEN-tools python package for more advanced users can be found at https://gitlab.ebi.ac.uk/cansu/centools_pycen.git. CEN-tools is currently available as a docker image which has all the functionalities of the website: ftp://ftp.ebi.ac.uk/pub/contrib/petsalaki/CEN-tools/. A detailed guide on using the docker image is provided in the appendix section. The website is currently being beta-tested and once stable will be shared through an updated version of this manuscript on bioRxiv. All data and networks will be available for download through the website and are available through the dockered app. All code and data are open source and freely available under a GPL license.

