## Supplementary figures for "CEN-tools: An integrative platform to identify the ‘contexts’ of essential genes"

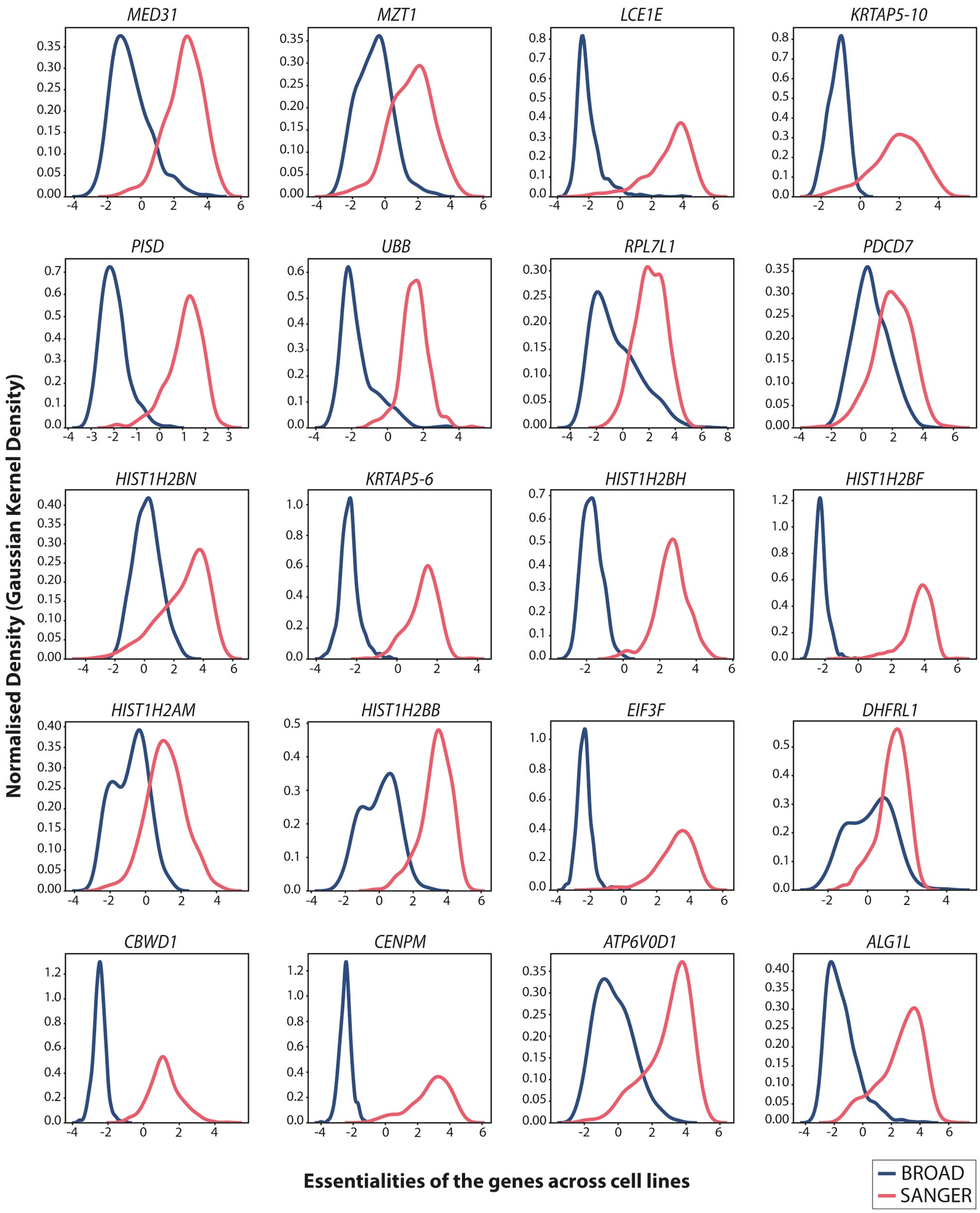


**Fig EV1. Project-specific discrepancy of essentiality profiles was observed for a number of genes.** Essentiality profiles of previously annotated genes from the ADaM pipeline (Behan et al. 2019), which were not identified as core essential genes from CEN-tools analysis pipeline. The profiles showed major discrepancies between the two projects. The ADaM pipeline utilised the essentiality screens from Project Score.

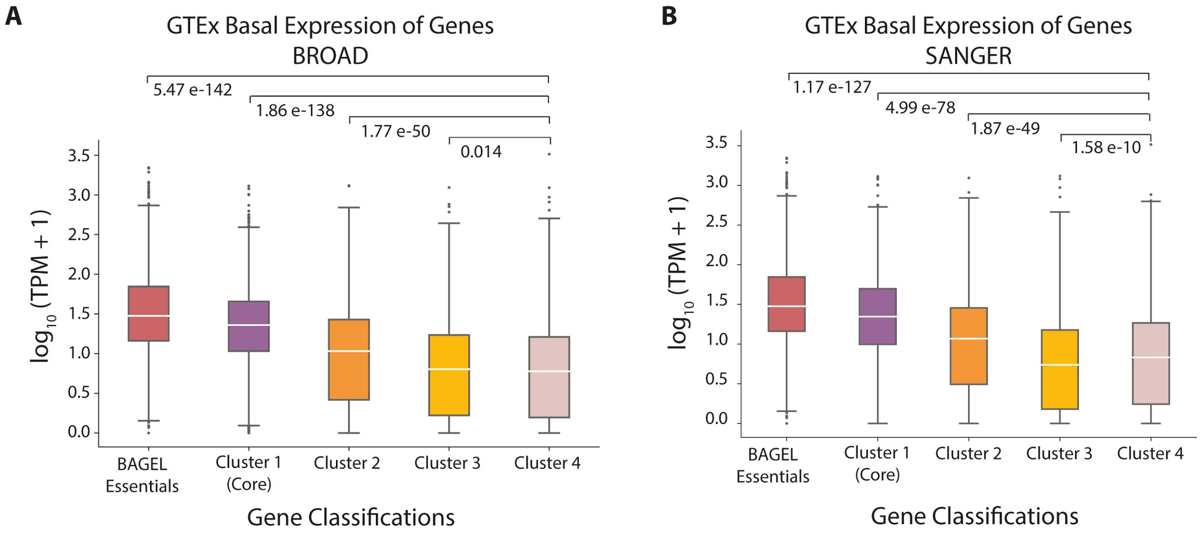


**Fig EV2. Core essential genes have higher basal expression in normal tissues than non-core genes.** Expression values were obtained from GTEx (Stranger et al. 2017). Expression values in the form of transcripts per million (TPM) with pseudocount of 1 are depicted for each of the four clusters identified from the core-analysis pipeline of CEN-tools for (A) BROAD project and (B) SANGER project. Clusters are based on essentiality probability distributions with cluster 1 designated as the core-cluster. All comparisons were performed with cluster 4 as it was the cluster with essentiality probability distribution skewed to 0, indicating non-essentiality (also see **Fig EV10**).

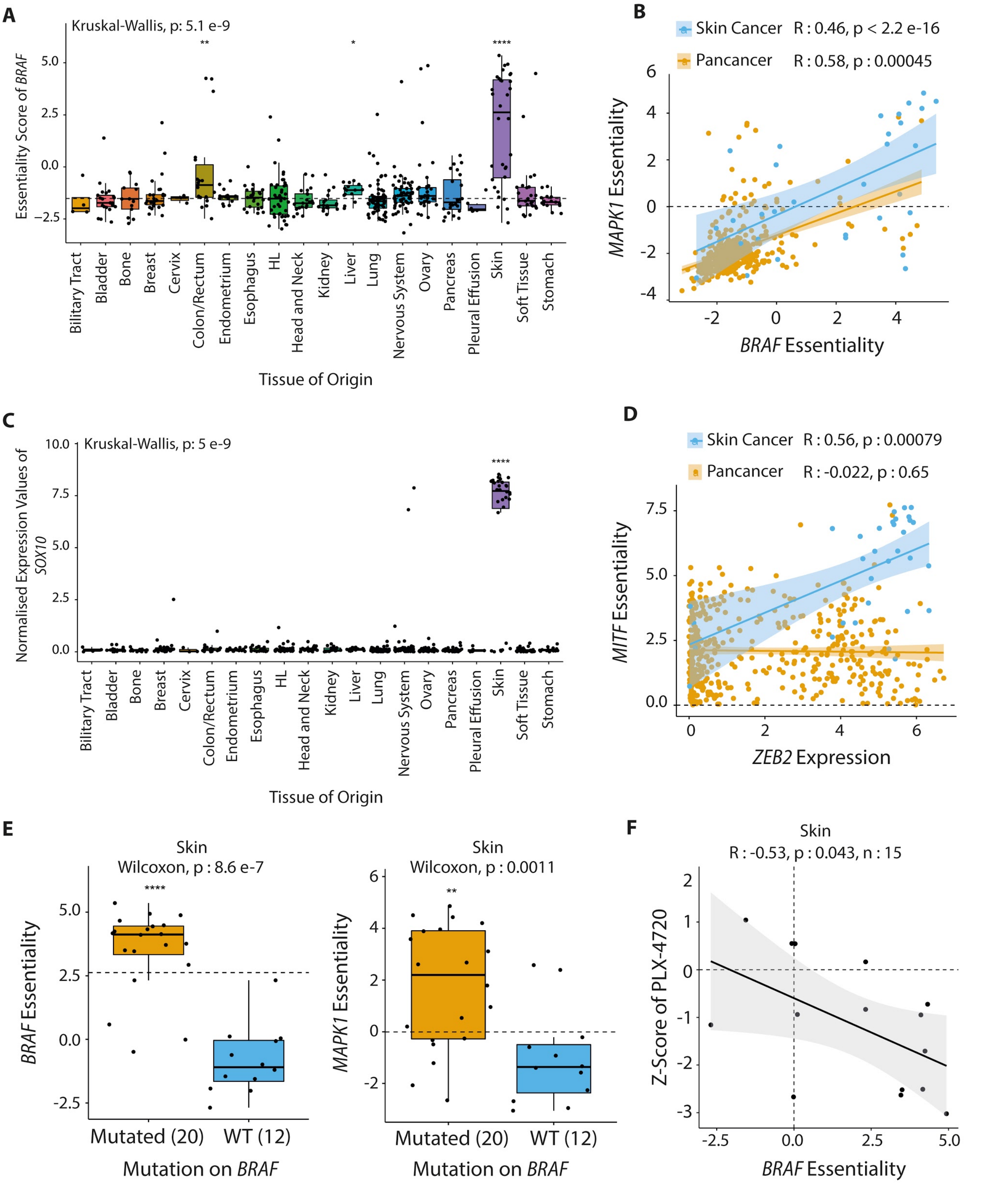


**Fig EV3. Representative plots from the CEN-tools website for predefined contexts.** (A) Tissue/cancer type-wide comparisons of *BRAF* gene essentiality, (B) essentiality correlations of *BRAF* and *MAPK1* genes in skin tissue compared to pancancer, (C) tissue/cancer type-wide comparisons of *SOX10* gene expression, (D) correlation between essentiality of *MITF* and expression of *ZEB2* in skin tissue compared to pancancer, (E) essentiality of *BRAF* and *MAPK1* in skin cell lines harbouring a *BRAF* hotspot mutation (BRAFV600E) compared to skin cells with WT *BRAF* and, (F) correlation between drug response to PLX-4720 (*BRAF*

inhibitor) and *BRAF* essentiality in melanoma cell lines. All plots show cell lines of the BROAD project.

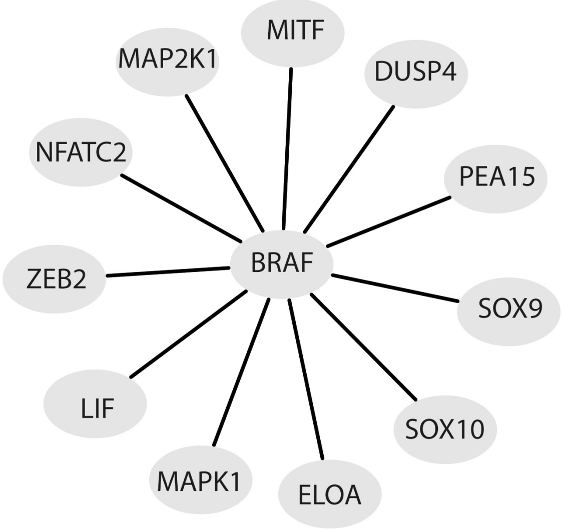


**Fig EV4.** The co-essentiality networks of *BRAF* obtained from the PICKLES web-server (Lenoir et al. 2018).

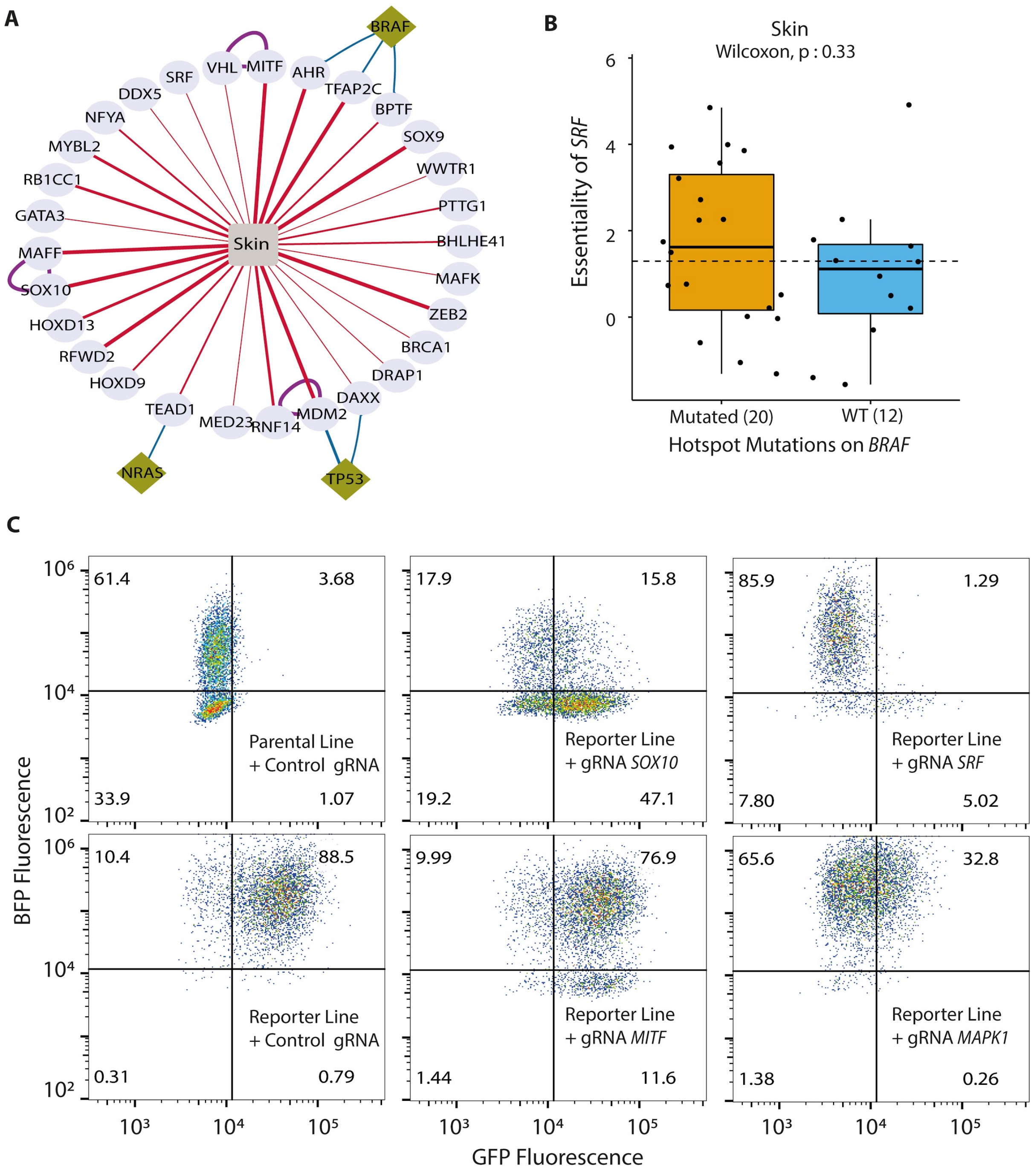


**Fig EV5. The utility of CEN-tools in identification of tissue-specific gene-gene relationship. (A)** The CEN of transcription factors in skin tissue. This CEN was generated from the BROAD dataset, which contains 32 cell lines from skin tissue. **(B)** The essentiality of *SRF* in skin tissue is not related to the *BRAF* mutational status of the skin cancer cell lines of the BROAD project. **(C)** Representative FACS plots depicting the effect of targeting denoted genes in the expression of GFP from the SRF-reporter construct.

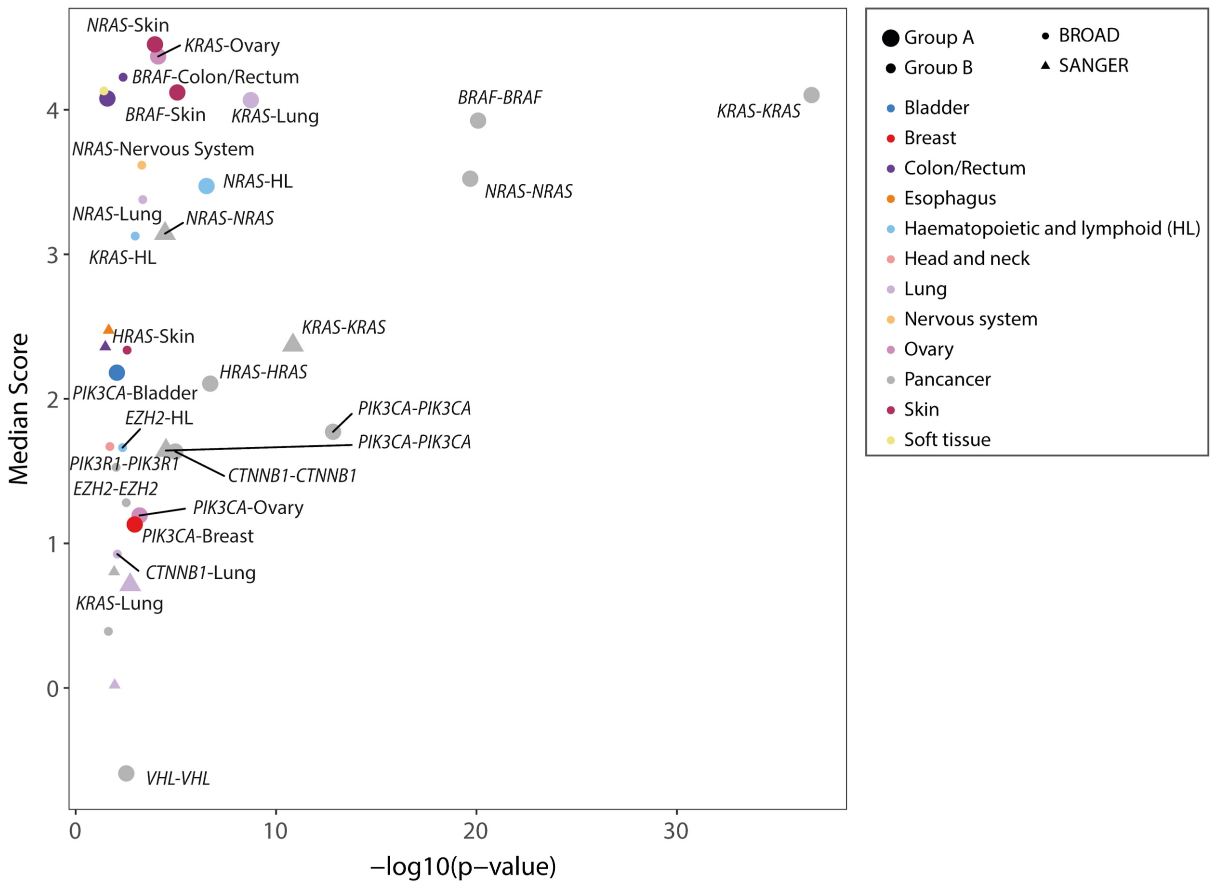


**Fig EV6. CEN-tools reveals mutation dependent vulnerabilities.** The essentiality of oncogenes and tumor suppressor genes in the context of their own mutations in pancancer and within tissue comparison. The 'Median Score' refers to the median of the scaled-Bayes essentiality score for cell lines from the indicated project for the indicated comparison. Group A and B refer to confidence of association with Group A being higher confidence in which number of samples/group was higher than 5, compared to higher than 3 for Group B comparisons.

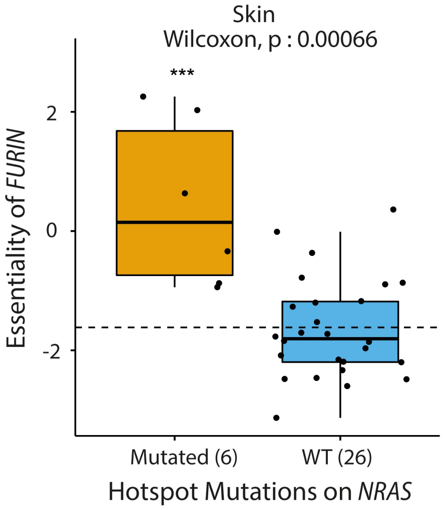


**Fig EV7. The essentiality of FURIN in *NRAS* mutant skin cell lines is significantly higher compared to that in *NRAS* WT melanoma cell lines.** Essentiality information from 32 skin cell lines of the BROAD project were used for this analysis.

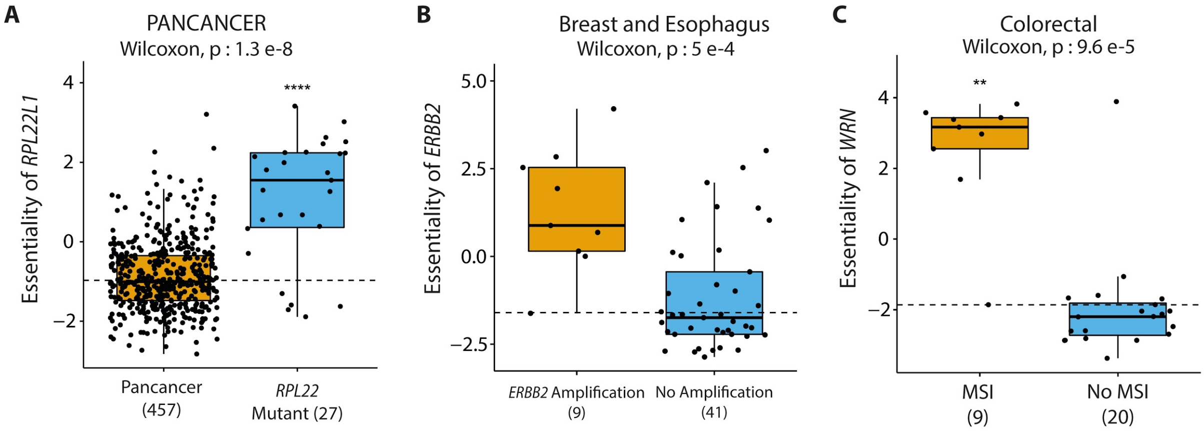


**Fig EV8. Examples demonstrating the utility of Cell Line Selector of CEN-tools in investigating essentiality in user defined contexts. (A)** Paralog dependency: Cells with mutation in *RPL22* show dependence on paralog *RPL22L1*. **(B)** Essentiality based on CNV status: Selection of cells with amplification of *ERBB2* gene in breast and esophagus tissues reveals increased dependency on *ERBB2* gene. **(C)** Essentiality based on microsatellite instability status (MSI): Colorectal cell lines with MSI show increased dependence on *WRN*. In all cases, cell lines in the defined context were selected from the Cell Line Selector application of CEN-tools. **(A)** and **(B)** show cell lines of the BROAD project and (C) shows cell lines of the SANGER project.

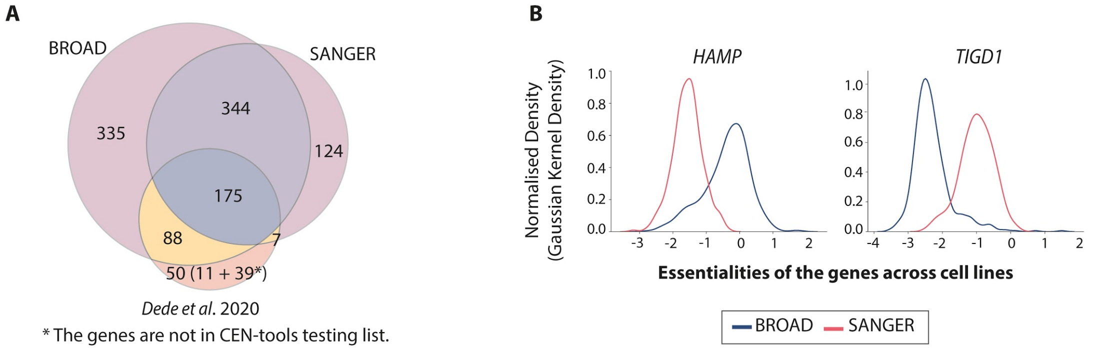


**Fig EV9. Comparison of CEN-tools core essential gene predictions with the predictions from *Dede et al*. 2020 (Dede et al. 2020) (A)** Venn diagram of the core predictions from CEN-tools using both BROAD and SANGER, from *Dede et al*. Only the overlap of the newly annotated core genes from *Dede et al* are shown. 39 genes from the *Dede et al.* analysis were not in the overlapping set of genes between SANGER and BROAD, hence were not present in the testing set of CEN-tools. **(B)** Essentiality distributions of *HAMP* and *TIGD1* as representative examples of genes predicted as core-essential genes from *Dede et al.*, but not from CEN-tools. The essentiality distribution profiles of these genes show inconsistencies between the SANGER and the BROAD projects.

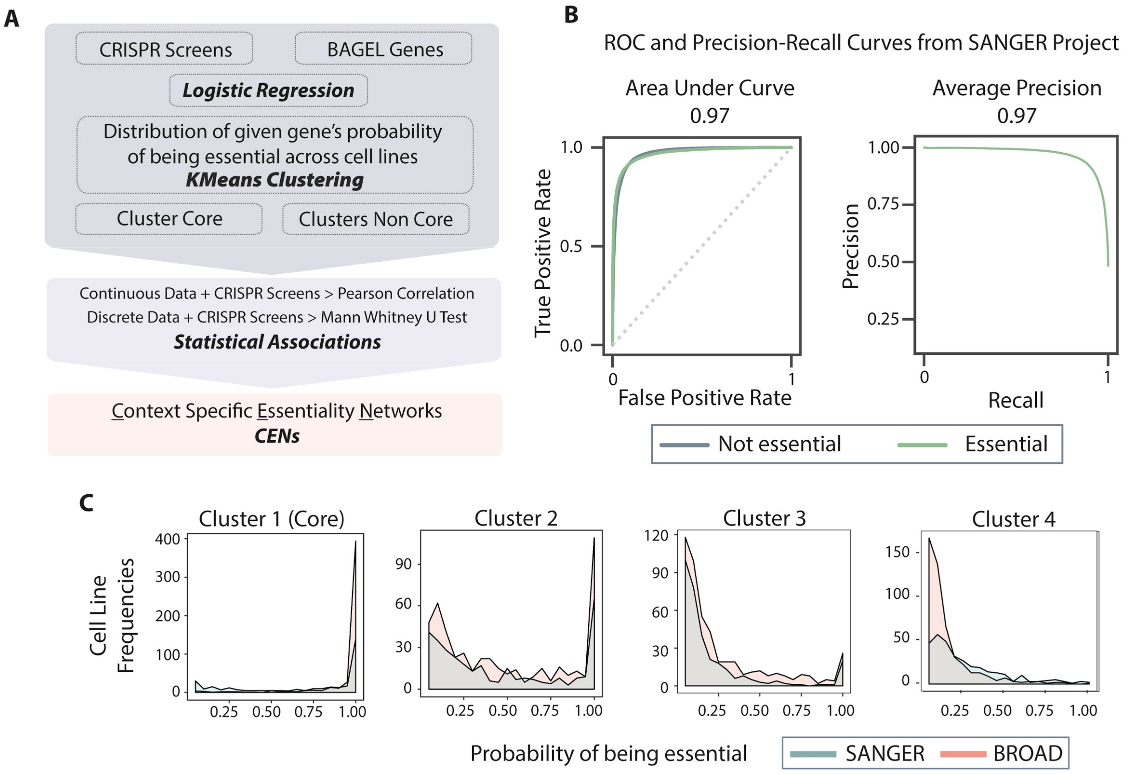


**Fig EV10. Workflow used for the identification of core essential genes in CEN-tools. (A)** The general workflow. **(B)** ROC and PR curves of the Logistic Regression (LR) algorithm. **(C)** Representative essentiality probability distributions from the four different clusters depicting the probability patterns.
